# Genetic transformation of micropropagated shoots of *Pinus radiata* D.Don

**DOI:** 10.1101/030080

**Authors:** Jan E Grant, Pauline A Cooper, Tracy M Dale

## Abstract

***Key Message*** *Agrobacterium tumefaciens* was used to transform radiata pine shoots and to efficiently produce stable genetically modified pine plants.

*Abstract* Micropropagated shoot explants from *Pinus radiata* D. Don were used to produce stable transgenic plants by *Agrobacterium tumefaciens-mediated* transformation. Using this method any genotype that can be micropropagated could produce stable transgenic lines. As over 80% of *P. radiata* genotypes tested can be micropropagated, this effectively means that any line chosen for superior characteristics could be transformed. There are well-established protocols for progressing such germplasm to field deployment. Here we used open and control pollinated seed lines and embryogenic clones. The method developed was faster than other methods previously developed using mature cotyledons. PCR positive shoots could be obtain within 6 months of *Agrobacterium* co-cultivation compared with 12 months for cotyledon methods. Transformed shoots were obtained using either kanamycin or geneticin as the selectable marker gene. Shoots were recovered from selection, were tested and were not chimeric, indicating that the selection pressure was optimal for this explant type. GFP was used as a vital marker, and the *bar* gene, (for resistance to the herbicide Buster^®^) was used to produce lines that could potentially be used in commercial application. As expected, a range of expression phenotypes were identified for both these reporter genes and the analyses for expression were relatively easy.

## Introduction

*Pinus radiata* D. Don (radiata pine, Monterey pine) is an important plantation forestry species in the Southern hemisphere, particularly in New Zealand, Australia and Chile. Pine wood and pulp is widely used in the construction and newsprint industries. Gains in plantation forest productivity have been made through advances in silvicultural practices and genetic improvement of planting stock by traditional plant breeding. As with other outcrossing forestry species, genetic engineering offers the opportunity to modify quality traits in a shorter time frame than traditional breeding.

Genetic modification of conifers has relied upon microprojectile bombardment and more recently, on *Agrobacterium tumefaciens–mediated* methods in combination with somatic embryogenesis from cultured immature zygotic embryos to produce transformed trees (*Larix kaempferi x L decidua*: Levée et al. (1997); *Pinus radiata;* Walter et al. (1998), Cerda et al. (2002); *Pinus strobus:* Levee et al. (1999); *Picea abies:* Wenck et al. (1999), Brukhin et al. (2000), Klimaszewska et al. (2001); *Picea glauca, P. mariana:* Klimaszewska et al. (2001). Cotyledon explants from mature *Pinus radiata* embryos have been successfully transformed with *Agrobacterium tumefaciens,* and transgenic plants have been produced from a range of *P. radiata* genotypes from open-pollinated and control-pollinated seed (Grant et al. 2004). For loblolly pine, *(Pinus taeda),* Tang et al. (2001) and Tang (2003) reported successful *Agrobacterium* transformation of mature embryos and for chir pine *(Pinus roxbughi)* Parasharami et al. (2006) used particle bombardment to transform mature embryos.

The major disadvantage for transformation of cultured embryogenic lines from immature zygotic embryos is that, generally very few genotypes can be successfully regenerated to somatic embryos and subsequently transformed by any methods. The use of mature embryos from seed, which are available year round, and the ability to regenerate adventitious shoots from the majority of genotypes can overcome the issue of limited genotypes. Gould et al. (2002) transformed shoot apices in loblolly pine with kanamycin selection at a ‘leaky or permissive’ level in the selection medium resulting in some chimerism of the recovered shoots.

In *P. radiata* the majority of genotypes are able to produce many adventitious shoots from the cotyledons (Aitken-Christie et al. 1988; Horgan and Aitken 1981; Smith 1986). Commercial clonal propagation of desirable genotypes by cuttings and shoots is well established in the New Zealand forestry industry. For *P. radiata* micropropagation through tissue culture is also established in commercial operation. Over 90% of genotypes of *P. radiata* used in the forestry industry are amenable to micropropagation (D. Adam, Rayonier NZ, pers comm.). For outcrossing trees with long generation times testing of genotypes for desirability is time consuming and ideally any introduced genetic modification would be in tested genotypes.

Here we describe a method for *Agrobacterium tumefaciens* genetic modification of *P. radiata* micropropagated shoots in tissue culture. Advantages of this method are the availability of the target tissue, which is not dependent on a narrow window for collection as for development of embryogenic cell lines. The lines that used are from different genotypes and ages and can be field tested genotypes. It is rapid and the efficiency is comparable with other reported methods of *P. radiata* transformation. This method will enable breeders to select from known genotypes and superior performers. Changes made to selected genotypes can be quite specific depending on the gene(s) chosen.

## Materials and methods

### Plant Material

Shoot cultures were derived from seeds (both open pollinated GF 17, GF 19 and control pollinated GF 26), somatic embryos (gift from Rayonier NZ Ltd) and 3 year old hedges. Shoot cultures were maintained in culture using the medium described in detail in Grant et al. (2007).

### *A. tumefaciens* and binary vectors

*Agrobacterium tumefaciens* strain KYRT1 (Torisky et al. 1997) containing one of the binary vectors described below was grown overnight in Luria broth (LB) supplemented with 5mM MES (2-(*N*-morpholino)ethanesulfonic acid), 100 mg/L streptomycin, 40 mg/mL acetosyringone, 10 mg/mL tetracycline. The next morning the *A. tumefaciens* culture was centrifuged at 4,000 rpm for 6 min. The supernatant was discarded and the pellet resuspended in fresh LB supplemented with 0.5% DMSO (dimethyl sulfoxide) to the required density – OD 550 nm = 0.35 - 0.40.

### Plasmids

Six different plasmids were used in this study with a range of promoter sequences driving the selectable marker and reporter genes. Two of the plasmids have the *bar* gene as this was of interest to the industry partners in this project.

1. pMP2482 (Quaedvlieg et al. 1998) was kindly gifted by Dr. Spaink (Netherlands). This plasmid contains a *uidA* gene fused to a *sgfpS65T* gene (Chiu et al. 1996) and both are under the control of an AMV enhancer element and a double CaMV 35S promoter. The backbone for this construct is pBINPLUS (van Engelen et al. 1995). The *nptII* gene is under control of a *nos* promoter and *nos* terminator and is located proximal to the right border.
2. pTGUS (Liew 1994), kindly gifted by Dr Oi Wah Liew, contains the *uidA* gene under the control of a TMV enhancer region and the CaMV 35S viral promoter (CaMV 35S) and an *ocs* terminator in the backbone of pGA643 (An et al. 1988). The *uidA* gene is located proximal to the left border region. The selectable marker gene *nptII* is under the control of a *nos* promoter and *nos* terminator.
3. p4CL (Hu et al. 1998) kindly gifted by Dr Chung-Jui Tsai (MIT, USA) contains the 4-Coumarate:CoA ligase 1 promoter from *Populus tremuloides* driving the GUS gene and a *nptII* gene for kanamycin selection with a *nos* promoter and *nos* terminator.
4. pSLJ1111 (Scofield et al. 1992), kindly gifted by Jonathon Jones, is a binary vector that contains the *uidA* gene under control of the T_R_2′ promoter. This promoter is a constitutively expressed and was isolated from the Ti plasmid of *Agrobacterium tumefaciens* (Velten et al. 1984). The *nptII* gene is under the control of a CaMV 35S promoter. This vector carries an *Ac* transposase gene also transcribed by a CaMV 35S promoter.
5. pLN27 (constructed by Janet White and gifted by Dr Kevin Davies, Plant & Food Research, New Zealand), contains the *bar* gene (coding for phosphinothricin acetyl transferase) under the control of the CaMV 35S promoter and a 5’7’ terminator in the backbone of pGA643 (An et al. 1988). The *bar* gene is located proximal to the left border region. The selectable marker gene *nptII* is under the control of a *nos* promoter and *nos* terminator.
6. pLUG, constructed by Dr Richard Weld, Lincoln University, New Zealand, contains a CaMV 35S promoter driving the *bar,* a second CaMV 35S promoter driving the *uidA* gene and a ubiquitin promoter driving the *npt*II gene on pCAMBIA 3301 +pRN2 (4Kb HindIII frag) T-DNA = 9025 bp.

### Shoot transformation

Adventitious shoot explants, for transformation experiments were from actively growing cultures that had been subcultured onto 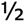 LP medium (Le Poivre medium as modified by Aitken-Christie et al. (1988)) with no growth regulators 3-4 weeks earlier. The apical stem was cut ~1cm from the top, the needles cut off and the stem cut longitudinally with a scapel that had been dipped in the *Agrobacterium* culture, to give 2 explants. Each explant was then dipped into the *Agrobacterium* culture and placed onto 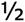 LP medium containing 5 mg/L 2,4-D and 20 mg/L acetosyringone. After 3 days explants were transferred to / LP medium containing 5 mg/L BA and 200 mg/L Timentin. After 7 days explants were transferred onto the same medium and after a further 7 days transferred to selection medium – 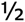 LP with 200 mg/L Timentin and either 10 mg/L kanamycin or 6 mg/L geneticin. Transfers were carried out fortnightly to selection medium for a minimum of 12 weeks. Shoots were then grown on as described in Grant et al. (2007).

### Transfer to soil

Transplanting shoots to glasshouse was carried out as in Grant et al. (2007).

*Agrobacterium* contamination

To determine if any PCR positive results were due to residual *Agrobacterium* infection in the plantlets, needles and stem pieces were grown in Luria Broth in the same conditions as in Grant et al. (2004).

### Molecular Analyses

#### DNA extraction

For small quantities of genomic DNA 100 mg (fresh weight) of *P. radiata* tissue was ground in liquid nitrogen using a plastic disposable pestle in eppendorf tubes and the DNeasy Plant Mini kit (Qiagen) extraction method, following the manufacturer’s instructions.

For larger quantities of genomic DNA 0.8-1.2 g (fresh weight) of *P. radiata* tissue was ground in liquid nitrogen using a pestle and mortar. Two methods of extraction were used:

1. DNeasy Plant Maxi kit (Qiagen), following manufacturer’s instructions,
2. A modified CTAB method (Doyle and Doyle 1990; Grant et al. 2004).

### Southern hybridisation

Southern hybridisation was carried out as described by Grant et al. (2004) and Dale (2004). Genomic DNA was digested with *Eco*R1 for pLN27 and pLUG, *Hind*III for pSLJ1111, pPMP 2482 and p4CL, which cut once in the T-DNA. In addition genomic lines transformed with pPMP2482 were cut with EcoR1 which cuts twice within the T-DNA and *Pst*1 which cuts three times.

### PCR Analyses

Each PCR was performed several times to ensure reproducibility. PCR’s to confirm the presence of the transgene included the selectable marker gene - *nptII*, and the gene(s) of interest - *uidA, gfp* and *bar.* For further confirmation spanning PCR, where primers were used in various combinations to amplify PCR products spanning the promoter, the terminator, the selectable marker and the gene(s) of interest was carried out. To ensure the result was not due to *Agrobacterium* contamination, PCR of the virulence gene *VirG* was carried out on all putative shoot transformants.

PCR analysis was carried out on several different DNA extractions from putatatively transformed shoots to check that the plants from one original shoot were not chimeric. For expression studies, DNA from at least 5 shoots was extracted separately and all 5 were tested by PCR for the selectable marker gene, the gene(s) of interest and *Agrobacterium* virulence genes.

### Primers used

The following primers were used in the illustrations in this paper.

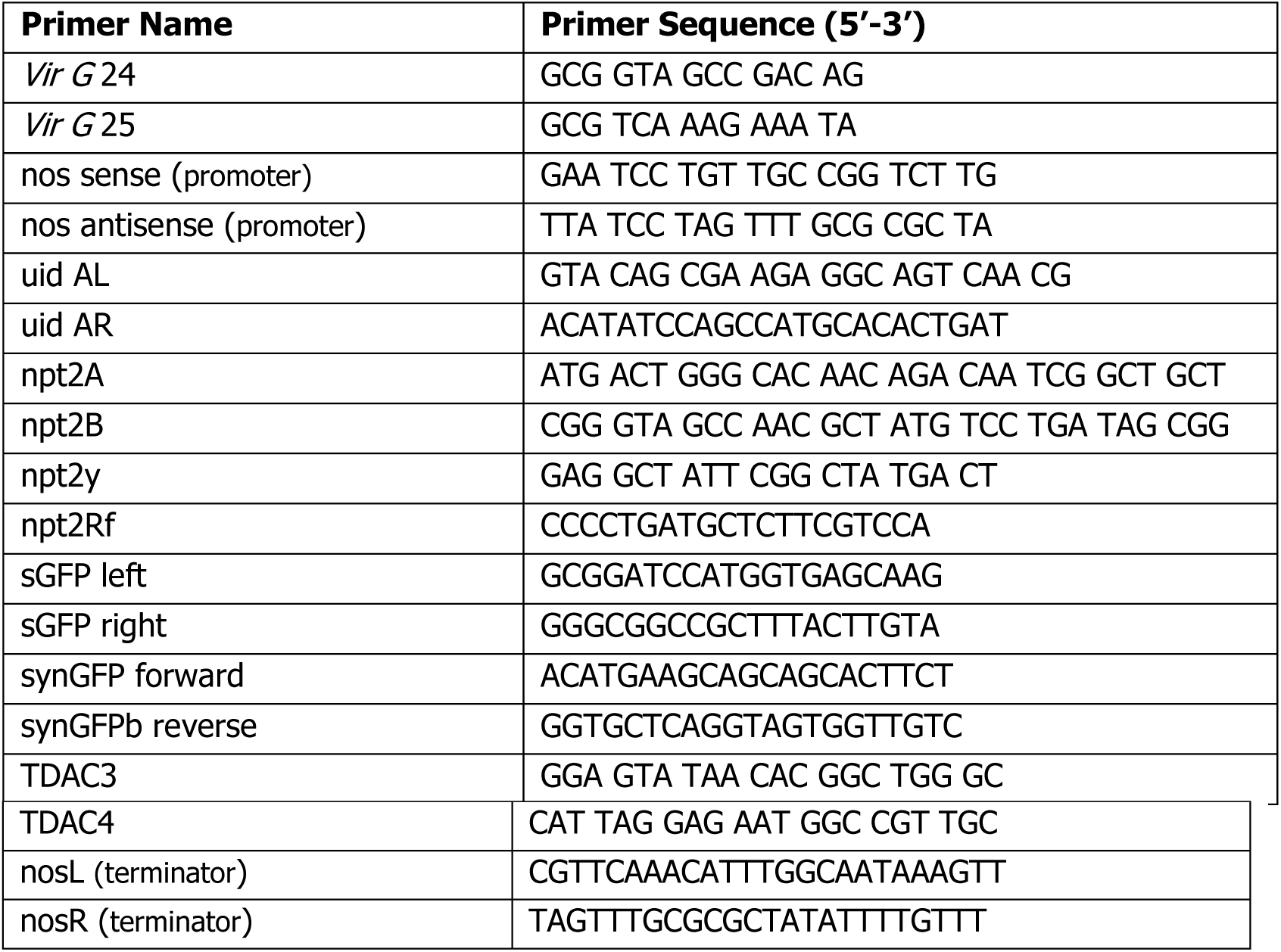

### Gene expression

#### Growing shoots on Buster^®^/Basta^®^

To test expression of the *bar,* putative transgenic shoots were grown on 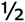 LP medium containing 5 mg/L phosphinothricin (PPT) if they were young shoots, or 8mg/L PPT for older shoots and for more stringent selection. Levels of PPT suitable for selection of transgenic shoots were previously determined by T Dale (unpublished data).

#### GFP

For expression studies, detection of GFP in needles was by fluorescence microscopy at 400x magnification. Young, actively growing needles from the tip of *P. radiata* shoots, were mounted in water and observed for fluorescence.

GFP fluorescence was quantitated for 7 independently transformed lines and 3 non-transformed controls by image analysis. Three needles were removed from the shoot tip of each shoot of the 7 transformed lines and 3 control lines. Epi-fluorescence microscopy at 10x magnification and identical procedures were used to capture the Red-Blue-Green channels, three images per needle were obtained. Within each image 5 standardized areas were measured for brightness of individual pixels from the green channel signal. The mean pixel brightness for each area was used for further data analysis by ANOVA. Data collection was done on different days using different needles to ensure reproducibility (for details see Dale 2004).

## Results

All constructs contained *npt*II so the explants could be selected on kanamycin or geneticin which had both proved to be successful for selecting transgenic *P. radiata* shoots from mature embryo explants (Grant et al. 2004). Both kanamycin and geneticin were successful for selecting PCR positive transformed shoots (Table 2) irrespective of the promoter sequence used to drive the *nptII* gene. The position of the *nptII* gene on the plasmid was not considered. Table 2 also shows that this method was successful with the range of different plasmids used and was irrespective of explant genotype.

**Table 2.** The selection agent, number of explants, genotypes and the range of plasmids used. All experiments produced PCR positive shoots. * PCR positive for at least 2 genes in the T-DNA and PCR negative for *Agrobacterium* contamination. ** Efficiency – Shoots from the same explant may or may not be identical. For the purpose of this efficiency estimate only one shoot from each explant is counted. *** embryogenic clones gifted by Rayonier NZ

| Experiment | genotypes | plasmid | No. explants | Selection | No. PCR positive shoots* | Efficiency** |
| --- | --- | --- | --- | --- | --- | --- |
| SU | GF17 & GF26 | pSLJ1111 | 335 | 10 mg/L kan or 6 mg/L gen | 13 | 3.9% |
| SW | GF19 | pSLJ1111 | 214 | 10 mg/L kan | 9 | 4.2% |
| SX2 | GF17 | p4CL | 160 | 10 mg/L kan | 8 | 5.0% |
| SX3 & SX4 | GF17 & GF19 | pPMP2482 | 225 | 10 mg/L kan | 8 | 3.5% |
| SY | embryogenic clones*** | pLN27 | 326 | 10 mg/L kan | 13 | 4.0% |
| TE | GF19 | pLUG | 193 | 10mg/L kan | 3 | 1.6% |
| Total |  |  | 1260 |  | 54 | 4.3% |

Fig 1 shows a summary of the steps to obtain transgenic shoots from the co-cultivation of pine shoot tip explants with *Agrobacterium tumefaciens* strain KYRT1. Fig 1a shows actively growing shoots ready for processing for transformation. After cocultivation for 3 days GUS expression spots were shown to be generally well spread along the cut surface of the shoot explants (Fig 1b). Non-transformed tissue would die and green shoots developed and grew until well established on selection medium (Fig 1c & d). This pattern was consistent for kanamycin and geneticin as the selection agent. Figure 1e shows transgenic plants in the greenhouse. Figure 1f shows a control needle with no GFP fluorescence only the red auto-fluorescence attributed to chorophyll. Fig 1g shows GFP fluorescence in a young, actively growing needle from a transgenic plantlet at 400X magnification. This sample shows green mesophyll cells indicating strong GFP expression along with the red auto-fluorescence from chorophyll. In needles from some transgenic plantlets where the GFP gene was present and the lines were low expressors, the GFP expression tended to be masked by the strong red auto-fluorescence of chlorophyll, although squashing the needles did release the fluorescing mesophyll cells.

**Fig. 1.**
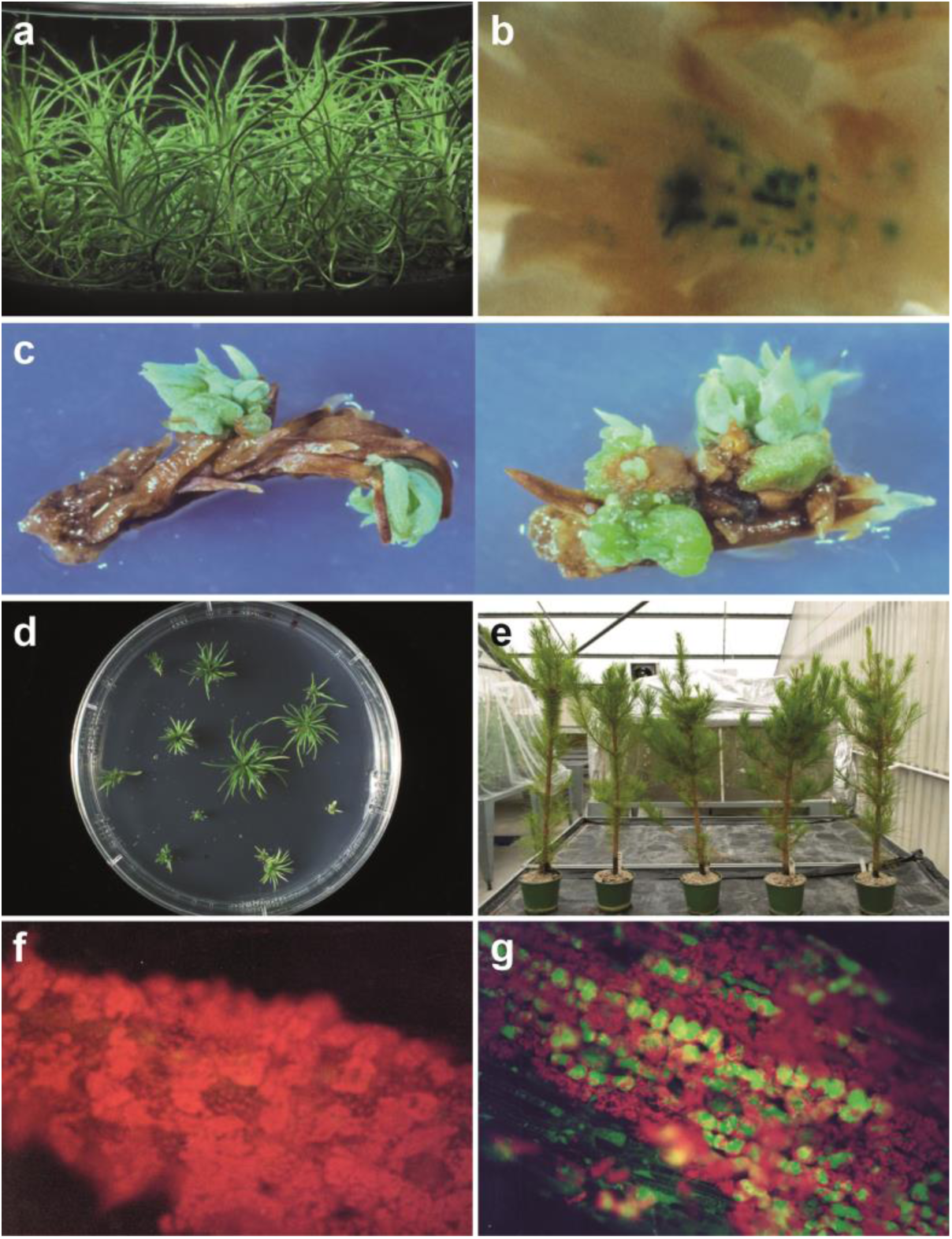
A summary of the production of a transgenic *P. radiata* line after *Agrobacterium* infection of shoot explants. a Micro-propagated *P.radiata* shoots ready for transformation. b Transient expression of the visual marker GUS (β-glucuronidase) in a *P.radiata* shoot explant, three days after cocultivation with *Agrobacterium* containing the plasmid construct pLUG. c Cocultivated shoot explants regenerating on selective medium containing 10mg/L kanamycin. d Shoots regenerated from explants after 18 weeks on selection medium. e Transgenic plants from 5 lines growing in the glasshouse. f A non-transformed *P.radiata* shoot shows the intense red auto-fluorescence of chlorophyll which is excited at the same wavelength as GFP. g GFP expression observed in a needle of a transformed *P.radiata* shoot – the red auto-fluorescence is due to chlorophyll which can mask the GFP expression.

The shoots that were identified as PCR positive in screening from regenerated shoots (Fig 2) were multiplied and a minimum of 5 different shoots were tested by PCR to ensure that chimerism was unlikely. This testing also ensured that the tissues used in expression studies and for Southern analyses were positive for the genes of interest. Further testing of needles and/or shoots in LB medium showed no evidence of latent *Agrobacterium* contamination. Lines were also checked to confirm integration by using PCR to span between the gene(s) of interest and/or with promoter and terminator sequences. The PCR’s in Fig 3 span between the marker gene and the terminator sequence in lines transformed with plasmid pMP2482. These lines were selected on kanamycin and, as well as the *nptII* gene, this plasmid has a GUS gene fused to a GFP gene. Figure 3a shows the GFP gene with the *nos* terminator with expected size of the sequence, 720bp, in the transformed lines the same as in the plasmid. However, figure 3b shows that the PCR with GUS gene and the *nos* terminator showed different sizes. The size from the plasmid is as expected at 1650bp, however, the size of the sequence in transformed lines is 1000bp indicating a deletion in this part of the T-DNA integration.

**Fig. 2.**
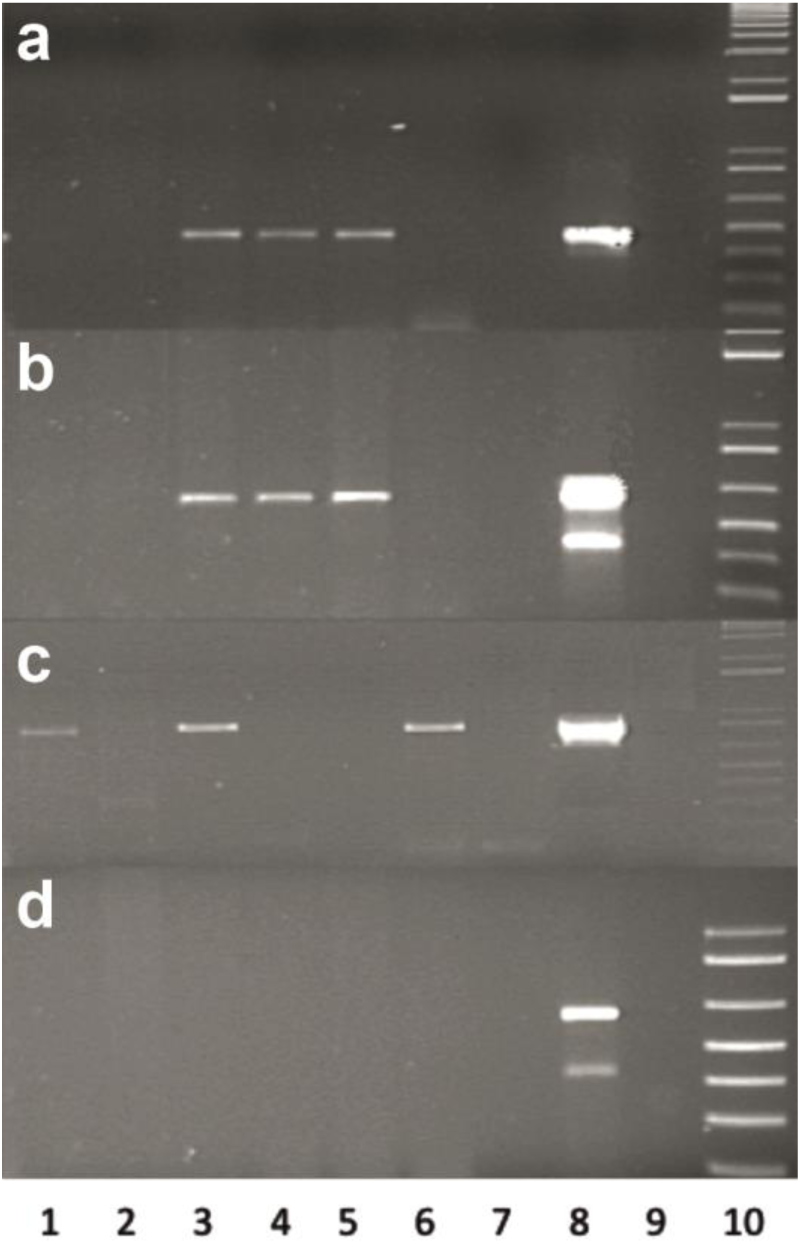
PCR of putative transgenic shoots for *uidA, nptll, AC, VirG* a PCR amplification of the *gus* gene using the primers uidAL and uidAR. The expected product size was 475bp. b PCR amplification of the *nptII* gene using the primers npt2a and npt2b. The expected product size was 837bp. c PCR amplification of the *AC* using the primers TDAC3 and TDAC4. The expected product size was 938 bp. d PCR amplification of the *virG* gene using the primers GMT24 and GMT25. The expected product size was 650 bp. Lane 1 –SU4.e1, lane 2 – SU4.e2, lane 3 – SW1.c2, lane 4 – SW1.f2, lane 5 – SW1.h2, lane 16 – SW1.h1, lane 17 – non-transformed control, lane 18 – pSJL1111 positive control, lane 19 – water blank and lane 20 – 1kb ladder (Invitrogen)

**Fig. 3.**
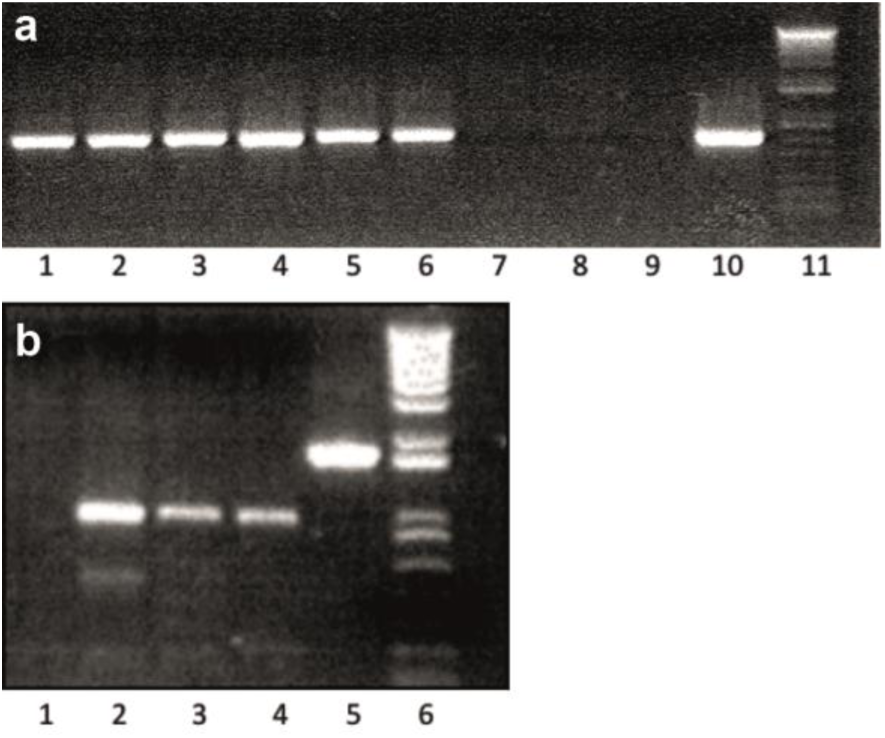
PCR spanning from the marker genes to the terminator sequence in lines with the plasmid pMP2482 to confirm integration of the introduced T-DNA. a PCR amplification from the *gfp* gene to the *nos* terminator in plasmid pMP2482 using primers synGFP and nosR. Lane 1-SX4.f1, lane 2-SX4.d2, lane 3-SX3.a3, lane 4-SX4.c1, lane 5-SX4.d1 lane 6-SX3.a1 lane 7-non-transformed control, lane 8-non-transformed control lane 9-water blank, lane 10-plasmid pMP2482, lane 11-1kb Plus ladder (Invitrogen) b PCR amplification from the *gus* gene to the *nos* terminator in plasmid pMP2482 using primers uidAL and nosR. Lane 1-water blank, lane 2-SX3.a3, lane 3-SX4.d2, lane 4-SX4.f1, lane 5-plasmid pMP2482, lane 6-1kb Plus ladder (Invitrogen)

Southern hybridization was carried out on PCR positive shoots using probes for the marker gene, other genes within the construct and promoter-gene combinations. In addition the blots were probed with a low copy number pine probe to check digestion and loading. As found in PCR’s not all introduced genes could be detected by Southern hybridization. Figure 4 shows a Southern hybridization for 9 putative transgenic pine shoots, that were shown to be positive for the *bar* gene using PCR. One line (lane 8) appears to be negative for Southern analysis and lanes 4 and 5 show the same hybridization pattern and as they come from the same original explant are likely to be identical. The lines are from co-cultivation of explants with 2 different plasmids pLN27 and pLUG each containing the *bar* gene driven by the CaMV 35S promoter. Digestion of the DNA with *EcoR1* cut once in the T-DNA in each of the plasmids pLN27 and pLUG.

**Fig. 4.**
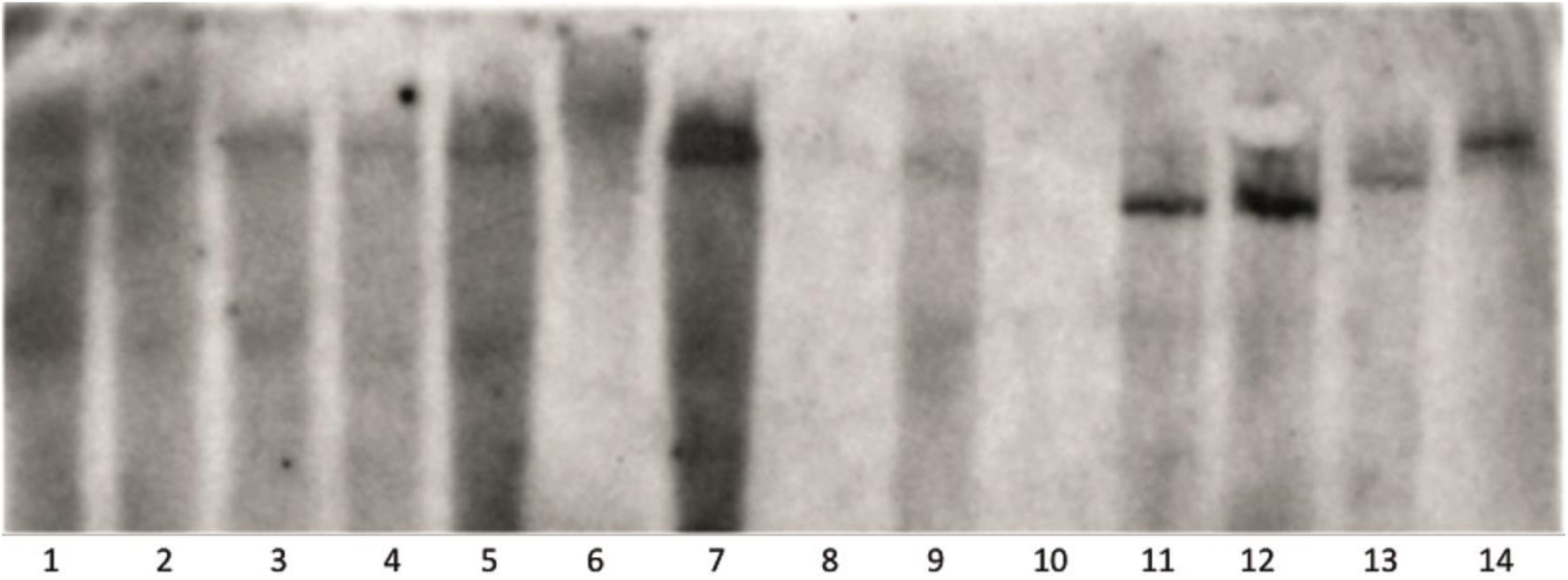
Southern hybridization of *Pinus radiata* digested with *Eco*RI and probed with *bar* gene primers Lane 1 –SY2.a1, lane 2 – SY3.a1, lane 3 – SY6.a1, lane 4 – SY6.c2, lane 5 – SY6.c3, lane 6 – SY7.a1, lane 7 – TD1.b1, lane 8 – TE3.a1, lane 9 – TE3.a2, lane 10 – non-transformed *P.radiata* control, lane 11 – control: *P.radiata* DNA + 10 pg pLN27 plasmid, lane 12 – control: *P.radiata* DNA + 20 pg pLN27, lane 13 – pLUG plasmid control: *P.radiata* DNA + 10 pg pLUG, lane 13 – control: *P.radiata* DNA + 20 pg pLUG plasmid

### Variation in gene expression

As expected there was a range of expression levels for the products coded by the inserted gene(s)

#### 1. Resistance to PPT

*P.radiata* shoots transformed with pLN27 and surviving on kanamycin selection were placed on / LP medium containing 5mg/L of PPT. Fig 5 demonstrates a single shoot surviving on 5mg/L PPT while the two other shoots show the typical browning of needles associated with PPT susceptibility.

**Fig. 5.**
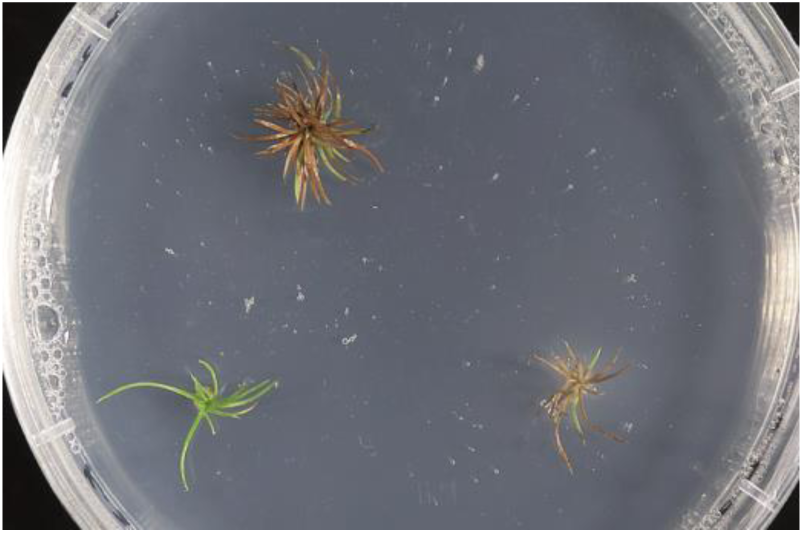
Selection of shoots on 5mg/L phosphinothricin (PPT) containing medium showing a single shoot surviving while the other 2 show browning of needles typical of PPT susceptibility As expected there was variation in resistance to PPT between the transgenic lines. Non-transformed control plants showed browning at 6 weeks and were completely dead at 12 weeks while the transgenic control plants on media without PPT remained healthy.

In Figure 6 Line SY6.c2 appeared to have the highest level of resistance to PPT and looked as healthy on media with 8 mg/L of ‘Buster’ (active ingredient 200g/L glufosinate-ammonium) as its control plants on media without PPT. Line TE3.c2 demonstrated a moderate to poor level of resistance to Buster. Other lines all showed varying degrees of resistance/tolerance to Buster containing media.

**Fig. 6.**
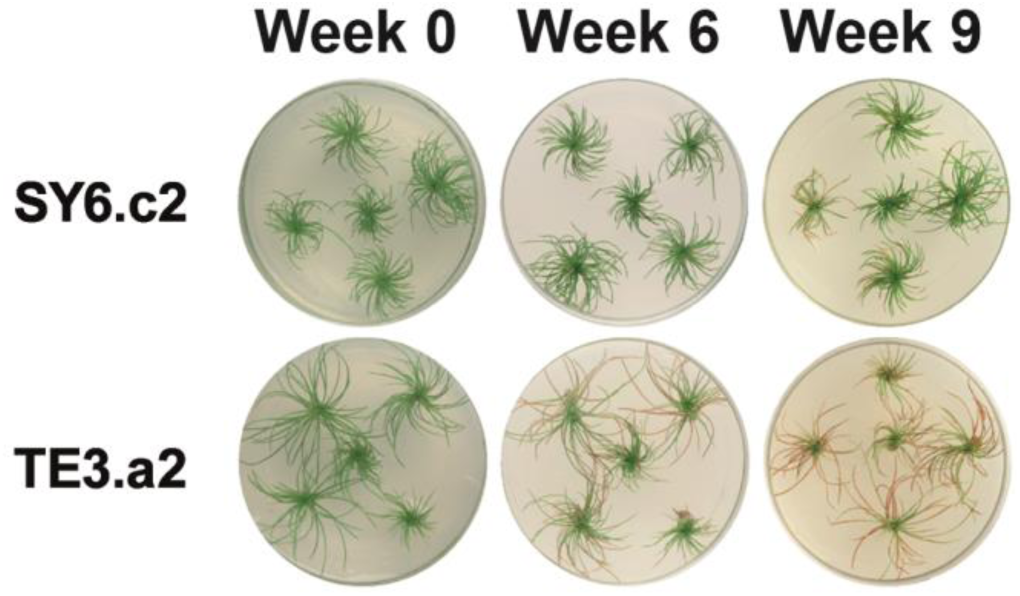
Different levels of susceptibility of transgenic lines containing the *bar* gene. Each line was grown on medium containing 8mg/L ‘Buster’ and photographed at 0, 6 and 9 weeks. Line SY6.c2 showed a good tolerance to ‘Buster’ while line TE3.a2 showed moderate to low tolerance

#### 2. Expression of GFP

GFP expression was determined by fluorescence microscopy. Red auto-fluorescence of chlorophyll masked GFP at lower levels of GFP expression but this could be eliminated by using an interference filter – Figs 1f & 1g shows negative and positive needles for GFP fluorescence without the use of an interference filter while Fig 7 shows the green fluorescence with the filter. Fig 7 shows differences in expression of needles in three independently transformed lines (Fig 7 a,b,c) along with a non-transformed control (Fig 7d). Weak auto fluorescence in the non-transformed control is due to cell wall components and other biochemical sources. That these lines were transgenic was confirmed by Southern analyses (Dale 2004 and unpublished data).

**Fig. 7.**
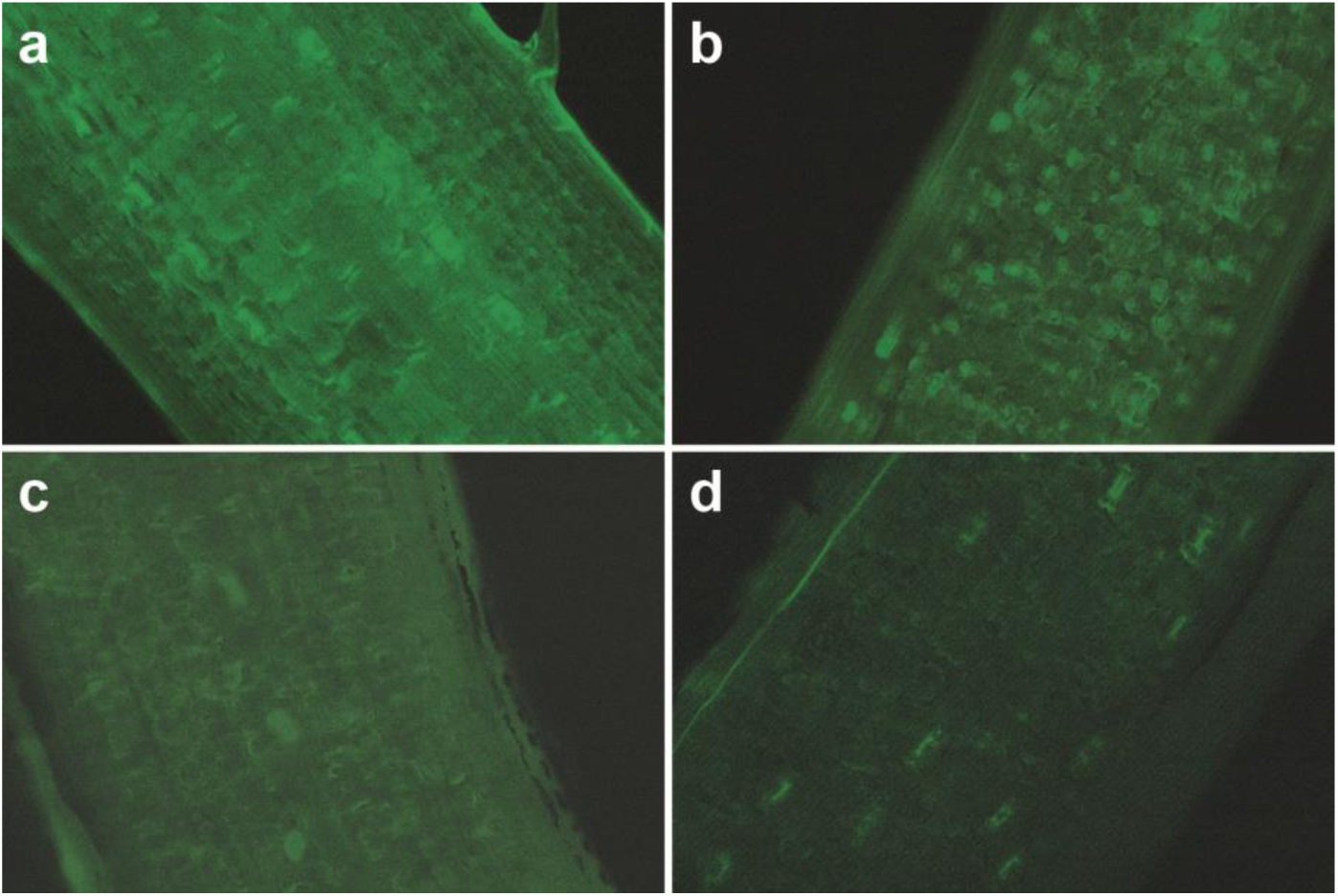
Detection of GFP fluorescence in *P.radiata* needles visualised by fluorescence microscopy. Using a *480/40nm* excitation filter and *510/40nm* emission filter observed under 400x magnification. a transgenic line SX4. f1, b transgenic line SX4.d1, c transgenic line SX4.h1 and d non-transformed control. Weak auto-fluorescence in the non-transformed control is due to cell wall components, phenolic compounds and other biochemical sources

Figure 8 shows the differences in green pixel intensity for seven transgenic lines and the 3 non-transformed controls. Within each needle the areas measured showed no differences in green pixel intensity. Samples from different days always had the same ranking *ie* line SX4.h1 always had the lowest intensity and SX4.f1 always had the highest. Three of the transgenic lines SX4.f1, SX4.d1 and SX4.h1 are the same as those shown in Figure 7 a, b and c respectively. Of the seven transgenic lines, all except SX4.h1 were readily distinguishable from the controls. Weak auto-fluorescence in the non-transformed control is due to cell wall components, phenolic compounds and other biochemical sources.

**Fig. 8.**
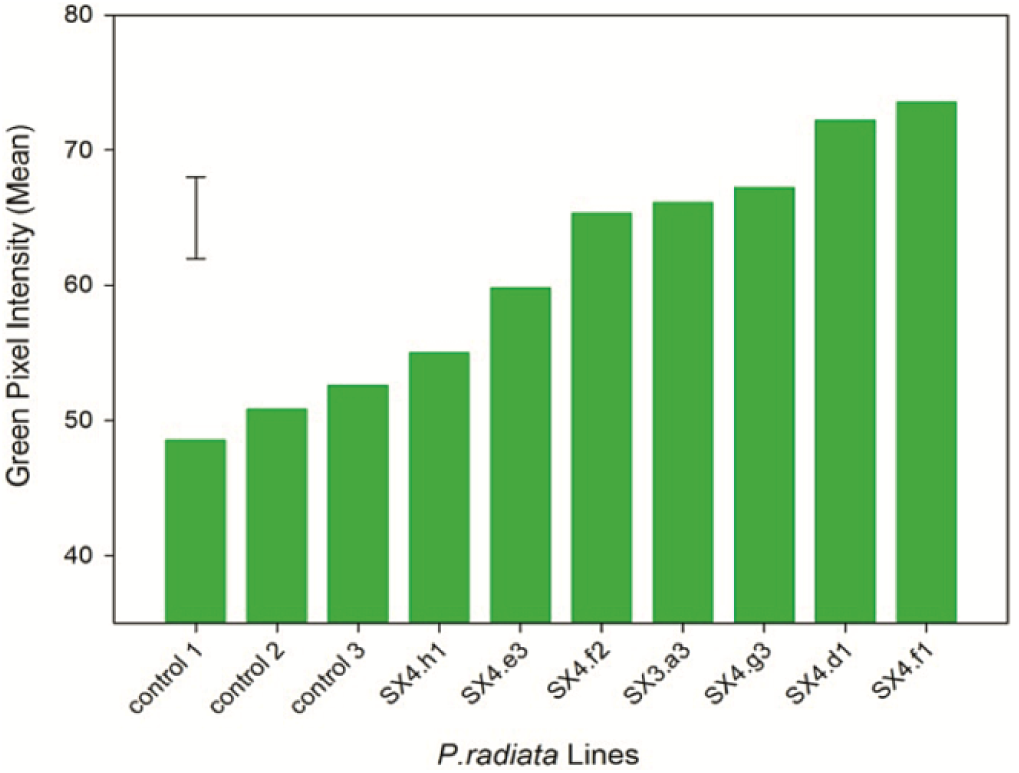
Independently transformed *P. radiata* lines had different levels of GFP fluorescence as determined by green pixel intensity. Significant differences in fluorescence were detected between lines (F=34.13, df=9,20, p=O.001). All transgenic lines except SX4.h1 were readily distinguished from the non-transformed controls

## Discussion

The ability to use adventitious shoots in pine transformation opens up opportunities to genetically engineer proven genotypes. Any line that can be introduced into tissue culture and produce adventitious shoots can be used in this method. We have shown shoots derived from open pollinated, control pollinated, 3 year old hedges, and embryogenic lines will produce transgenic plants. It is anticipated that genotypes that are rejuvenated in tissue culture or, by grafting and then introduced into tissue culture, could be transformed using this method. From co-cultivation of the explants with *Agrobacterium* to PCR positive transgenic shoots takes approximately 6 months. The transformation efficiency (number of explants producing one PCR positive shoot), at more than 4%, is similar to or better than other ‘recalcitrant’ species – *eg* apple up to 4.6% (Yao et al. 1995; Yao et al. 2013); onions up to 2.7% (Eady et al. 2000; Zheng et al. 2001) and for peas 0.8-13.4% depending on genotype (Grant and Cooper 2006; Grant et al. 1998). In comparison, explants from mature cotyledons and from somatic embryos generation of transgenic shoots takes at least 12 months (Charity et al. 2002; Grant et al. 2004) with transformation efficiency of 1.7%. Therefore the method presented here is much quicker and more efficient.

The promoter combinations with the marker and with the gene(s) of interest, their position on the T-DNA and T-DNA size did not appear to highlight a particular construct that was better or worse than the others. All constructs used produced some positive transgenic lines. This study again highlighted the difficulty of Southern analyses for large genome conifers when detecting single or low-copy number inserts (Charity et al. 2002; Grant et al. 2004). Southern analyses showed the majority of T-DNA insertions were single or low copy number and transgenic *P. radiata* lines that were highly expressing the gene of interest could be identified when using *nptII* as the selectable marker. Alignment of Southern hybridization reults to PCR results is imperfect although we found greater concordance when spanning PCR was used along within gene PCR. However we did find a large number of our transgenic lines with ‘non-classical’ (right border to left border) integration patterns (Dale 2004 and unpublished; Grant et al. 2004). Spanning PCR was used to confirm integration between genes and the various coding sequences. Such PCR’s indicated how the T-DNA was integrated. A curious feature of the spanning PCR pictured here was that all the lines shown had a similar deletion in the T-DNA although the shoots were from independent explants and/or separate experiments. Integration patterns in transgenic pine from both cotyledon explants and shoot explants have been further investigated using TAIL PCR (Liu et al. 1995) by Dale (2004) who showed that integration of the T-DNA included co-integration of plasmid backbone sequences and truncated or duplicated T-DNA sequences. Many authors using many different plant species and plasmids have since shown that right border to left border integration of T-DNA is not the normal occurrence (*e.g.* in barley, Bartlett et al. (2014); in apple, Yao et al. (2013); in *Arabidopsis,* Forsbach et al. (2003)). Detailed of results from both cotyledon and shoot integration patterns is part of a separate study (T Dale, pers comm).

The advantage of using GFP as a marker gene is that it does not require exogenous substrates or cofactors. The GFP sequence used here is a synthetic GFP with threonine at position 65 rather than serine which is in the native protein. Chiu et al. (1996) showed that this modified GFP protein showed greater expression in plant cells. Highly expressing lines can be easily determined by fluorescence microscopy in needles of forest trees (Tang and Newton 2005; Tian et al. 1999). In this study we developed a method that quantified the difference in GFP expression between lines as variation in green pixel intensity. The green pixel intensity was consistent within shoots of the same line and different between independent lines and the wildtype control. The lines showed expected differences in expression from highly expressing lines to lines where expression was almost the same as the non-transformed lines. Quantification of gene expression is important when choosing optimal expressing lines to go forward to further breeding and/or manipulation. This is especially pertinent for long lived forest trees as gene expression could be monitored over the life with its many changes in phases of growth and its environment. The method described for the quantitation of GFP in pines is relatively simple and non-destructive and was shown to give reliable expression information.

## Summary

The advantages of using adventitious shoots as explants includes:

- integration of genetically modified lines into well-established and effective protocols for progressing the germplasm to field deployment.
- the selectable maker *-nptII* is efficient to select transgenic lines and the GFP reporter and phosphinothricin (Buster) as marker genes confirmed the easy identification of highly expressing transformed genotypes.
- the use of a wide range of genotypes and selected specific genotypes, irrespective of age, as long as the line that can be propagated *in vitro*
- the opportunity to investigate mature tree characters in a short time frame *(cf* SE based systems).

## Author contribution statement

JG designed the research. JG, PC and TD carried out the plant transformation experiments. TD and PC carried out molecular biology experiments. TD conducted expression studies. JG wrote the paper.

## Acknowledgements

This research was supported by GEENZ Ltd, Rayonier NZ Ltd, Ernslaw One, Wenita Forest Products, New Zealand Foundation for Research, Science and Technology and a Bright Futures Enterprise Scholarship to TMD. We thank Lynn Thomson for technical support.

